# SBMate: A Framework for Evaluating Quality of Annotations in Systems Biology Models

**DOI:** 10.1101/2021.10.09.463757

**Authors:** Woosub Shin, Joseph L. Hellerstein, Yuda Munarko, Maxwell L. Neal, David P. Nickerson, Anand K. Rampadarath, Herbert M. Sauro, John H. Gennari

## Abstract

The interests in repurposing and reusing systems biology models have been growing in recent years. Semantic annotations play an important role for this, as they provide crucial information on the meanings and functions of models. However, there are a limited number of tools that evaluate the existence or quality of such annotations. In this paper, we introduce SBMate, a python package that would serve as a framework for evaluating the quality of annotations in systems biology models. Three default metrics are provided: coverage, consistency, and specificity. Coverage checks whether annotations exist in a model. Consistency tests if the annotations are appropriate for the given model element. Finally, specificity represents how detailed the annotations are. We analyzed 1,000 curated models from the BioModels repository using the three metrics and discussed the results. Additional metrics can be easily added to extend the current version of SBMate.

## 1 Introduction

The number of systems biology models has been rapidly increasing, thanks to the growing interest in using models to obtain biological insights and the improving software and hardware environments [6]. There are now several repositories that store curated models, such as BioModels and BiGG [6, 14]. BioModels has over 1,000 curated models, each of which contains on average about 30 reactions, derived from the literature available to users; BiGG currently has 108 genome-scale metabolic models, where the average number of reactions per model is over 2,000. These models are stored in the standard format SBML (systems biology markup language), which is a formalized machine readable description of biochemical reaction models [1]. The SBML standard allows modelers to add additional information to clarify the meaning of various terms in a model. Such information is referred to as model annotations. BioModels and BiGG are heavily annotated.

Repurposing publicly available models has become an intense interest among the modeling community in recent years; for example, model authors may want to identify and retrieve certain models that are relevant to their project, and reuse all or parts of such models. Semantic annotations can play a crucial role in this process, as they link metadata to controlled knowledge resources, allowing software tools to automatically recognize when different models and model elements represent the same or similar biological features [4, 13].

There exist a few studies that used semantic annotations to characterize and repurpose systems biology models. `semanticSBML`, a software package that merges multiple SBML models, evaluated and compared annotations to determine the similarity between models [5, 17]. `SBML Reaction Finder` searches reactions based on the user’s query and associated Gene Ontology (GO) terms [10]. Alm et al. proposed a similar approach, this time to extract SBML models that were relevant to specific biological themes (e.g., cell cycle or apoptosis), based on GO, CHEBI and SBO terms [2].

The success of these studies would depend largely on the quality of annotations and the correctness of them. So far, we have found only a limited number of previous studies on this. `semanticSBML` checks and reports to the user the model elements that were not annotated [7]. `MEMOTE`, a test suite for genome-scale metabolic models, included a section that examined whether reactions, species, and genes of a model were annotated, and whether such annotations were compliant with the community standards, i.e., minimum information required in annotation of models (MIRIAM) [8, 9]. Still, we lack a stand-alone package on testing annotations, with representative measures on the overall quality of model-level annotations in various aspects.

In our study, we have investigated approximately 1,000 curated models from the BioModels repository, which contained a total 30,094 reactions and 23,255 chemical species, and found various issues with the annotations provided in the models. Below describes common types of such issues:

- No annotation; BIOMD0000000608 had 52 species and 140 reactions, but none of them was annotated. In fact, among the 1,000 BioModels we examined, 9,502 out of a total 30,094 reactions did not have any annotation.
- Obsolete identifiers; in BIOMD0000000012, GO:0005623 (obsolete term for cell) were used to describe `cell` (compartment).
- Inappropriate identifiers; in BIOMD0000000356, reaction `v4f` had GO:0005942 (phosphatidylinositol 3-kinase complex), which describes an enzyme rather than a reaction itself.
- Insufficient information; in BIOMD0000000294, all reactions from `r1` to `r12` were annotated with SBO:0000375 (process), which, although correct, is too general and does not contain much information on the properties of the reactions.

The above issues could guide us into a few principles regarding how high-quality annotations would be like within a model. First of all, annotations should exist in order to be used or to be evaluated. Next, even if annotations exist, the terms should be ‘appropriate’, in the sense that the terms should come from collections of actively used terms, and also should match the characteristics of the object that it describes. Finally, it would be desirable for the terms to provide detailed information; for example, we would like the annotations contain more information than just *a reaction* or *a molecule*.

Based on these principles, in this paper, we provide three metrics, namely, **coverage, consistency**, and **specificity**, to evaluate and represent the quality of semantic annotations in systems biology models, especially those in the SBML format. Coverage measures whether a model component was annotated. Consistency indicates whether the annotation was appropriate given the type of model component. Finally, specificity examines how precise the annotation is. These measures will be described in more detail in the methods section. We implemented the three metrics in a convenient Python package called `SBMate` (Systems Biology Model AnnoTation Evaluator), which can be easily extended by the users as new metrics are developed.

We have two goals in our work: (a) first, to suggest the three metrics to evaluate the current state of model annotations, and (b) second, to introduce a model annotation testing framework that can be easily modified and extended by the users. Our results will be beneficial to the modelers who are seeking to appropriately annotate their models, and to those who want to validate and reuse already published models.

## 2 Methods

### 2.1 Scope of Analyses

#### 2.1.1 Annotations in Curated BioModels

We conducted an analysis of the annotations used in the approximately 1,000 curated ODE models in the BioModels repository.

In an SBML model, annotations can be added to various constructs, i.e., components of a model, represented as subclasses such as `libsbml.Reaction` and `libsbml.Species`. The full list of such constructs can be obtained by `getListOfAllElements()`, a libsbml method. In the following text, we will call such components **model elements**. In this study, we examined annotations in only four model elements; `libsbml.Model`, `libsbml.Reaction`, `libsbml.Species`, and `libsbml.Compartment`. We ignored the other components such as `libsbml.KineticLaw` and `libsbml.AssignmentRule`. The reason was that the four selected elements were most frequently annotated; more then 500 models had annotations for each of the four element types, either by the `sbo_term` attribute or the `getAnnotationString()` method. The other elements were less annotated in the current BioModels repository; for example, only 40 out of total 1,000 curated models provided an sbo term for `libsbml.KineticLaw`; none of them provided string annotation for kinetic laws. Figure 1 shows the number of models annotated per each major model element type. It can be easily seen that most models have annotations for the four model elements, while they have much fewer annotations for other element types. From now on, we call the instances of these four types of elements *annotatable*.

**Figure 1:**
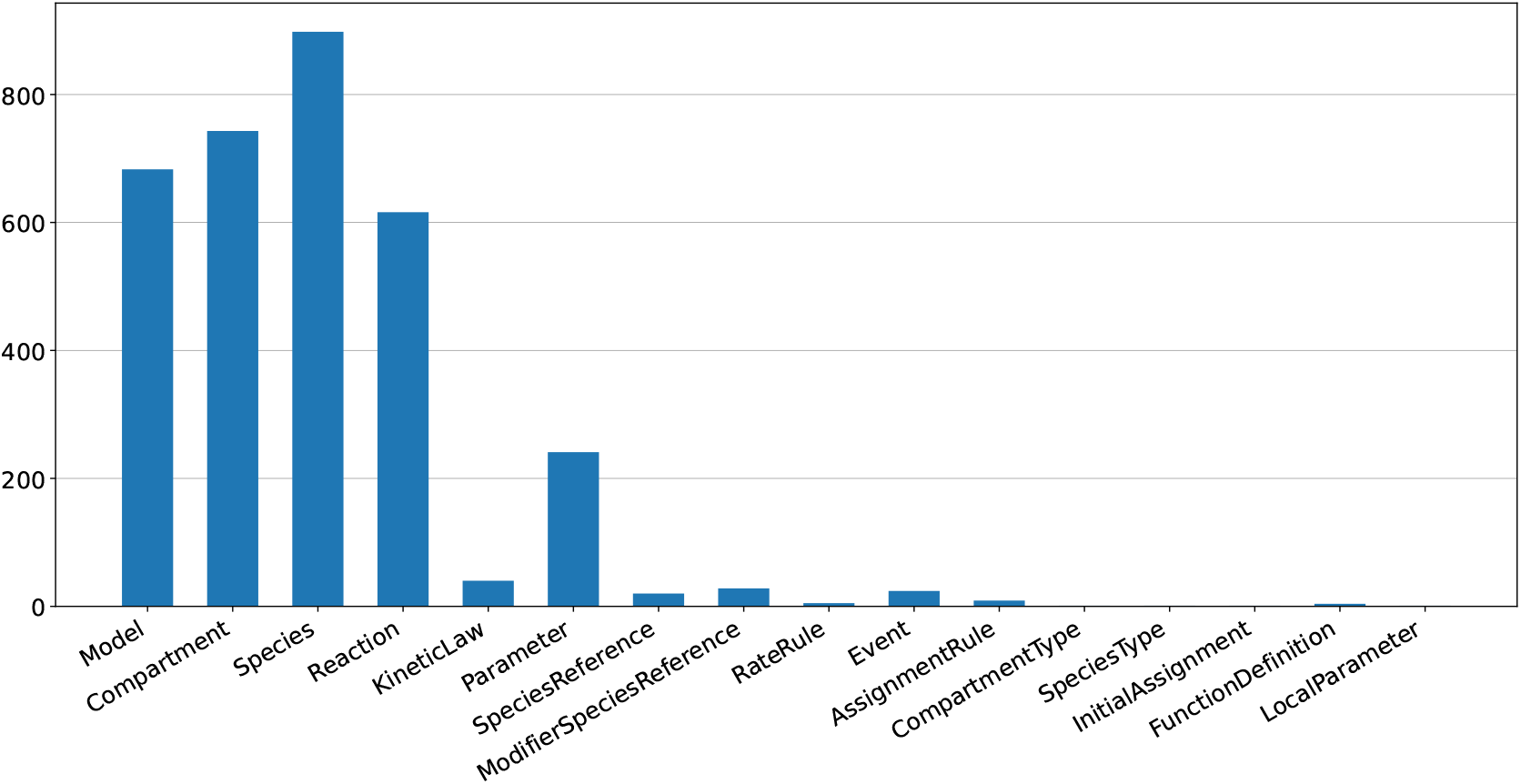
The number of annotated models for each model element type. Compartments, models, reactions, and species were the most well-annotated by modelers compare to the others.

An annotatable model element is *annotated* if its `sbo_term` attribute is an integer other than −1 (−1 means not provided) or its `getAnnotationString()` method returns at least one URI under the `bqbiol:is` or `bqbiol:isVersionOf` tag.

#### 2.1.2 Knowledge Resources

As in model elements, we limited our study to five knowledge resources out of many existing ones: GO, SBO, CHEBI, KEGG, and UNIPROT. One reason was that the five knowledge resources were well-suited to describe the four model elements. For example, `libsbml.Reaction` could be appropriately described as a term related to GO biological process (GO:0008150) or GO molecular function (GO:0003674). Similarly, a `libsbml.Species` could be a protein complex (e.g., CAF-1 complex, GO:0033186, or its subunit B, UNIPROT:Q13112) or a molecule (e.g., glycogen, CHEBI:28087), which could be annotated with a GO or UNIPROT or a CHEBI identifier. The second reason was that the five knowledge resources were the most frequently used in the BioModels repository. Figure 2 describes the proportion of knowledge resources in the BioModels repository, and the five ontology systems would count for about 85% of all annotations. The characteristics of the knowledge resources and their naming conventions are summarized in Table 1.

**Table 1:**
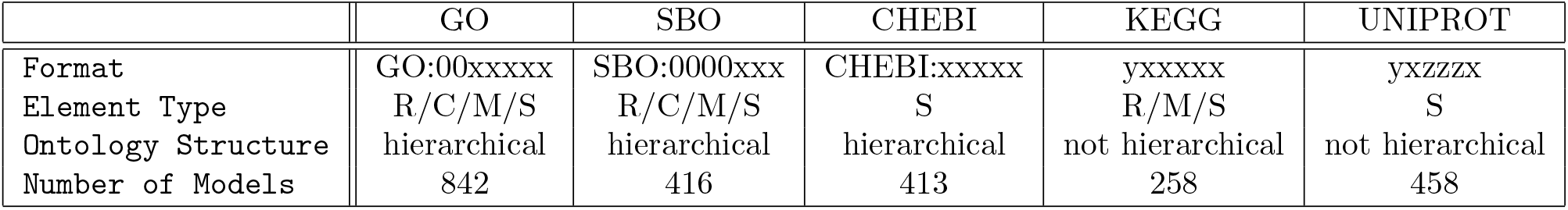
Summary of five knowledge resources. `R`: reactions, `C`: compartments, `M`: models, `S`: species, `y`: an arbitrary letter, `x`: an arbitrary number, `z`: an arbitrary letter or number.

**Figure 2:**
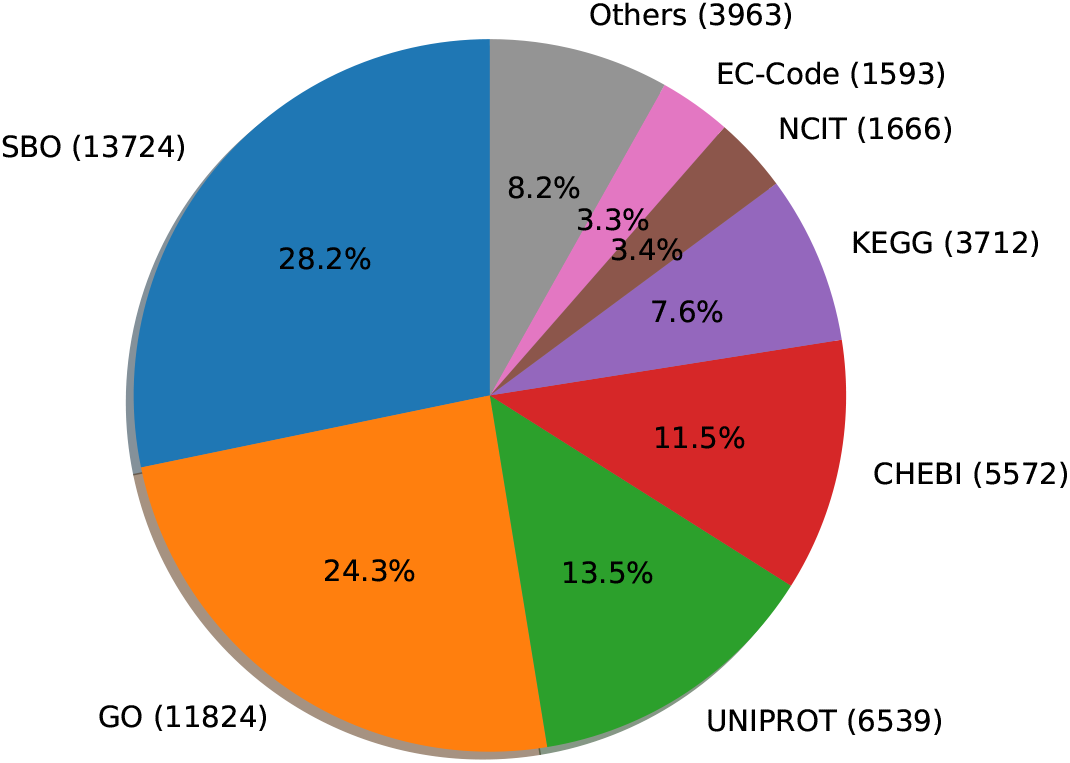
Distribution of knowledge resources of the four element types, obtained from 1,000 BioModels. Only seven most frequently used knowledge resources were specified. Annotations were collected either by the `sbo_term` attribute or the `getAnnotationString()` method. In annotation strings, those with either `bqbiol:is` or `bqbiol:isVersionOf` tag were selected. SBO and GO consisted more than 50% of all annotations found. In the following analyses, we used the five most common ontologies, i.e., SBO, GO, UNIPROT, CHEBI, and KEGG.

GO, SBO, and CHEBI terms are hierarchically organized and can establish a directed acyclic graph (DAG), where certain terms function as roots and other terms as children of others. For example, GO has three root terms: GO:0003674 (molecular function), GO:0008150 (biological process), and GO:0005575 (cellular component). Every GO term has one of them as its root. Similarly, all of SBO terms are descendants of SBO:0000000 (systems biology representation), and every CHEBI term has either CHEBI:24431 (chemical entity) or CHEBI:50906 (role) as its root. In this study, we ignored descendants of CHEBI:50906, as they describe certain biochemical functions rather than chemical species, for example, CHEBI:59132 (antigen) or CHEBI:39317 (growth regulator), and account for only about one percent of total CHEBI terms.

Compared to the three knowledge resources above, KEGG and UNIPROT describe each chemical species or chemical reactions and their terms are not hierarchically organized; for example, R00259 is a KEGG reaction defined as:

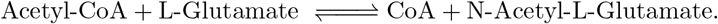

Therefore, we consider each valid KEGG or UNIPROT term as equivalent to a leaf of the hierarchical ontology graph.

### 2.2 Metrics: Coverage, Consistency, and Specificity

Here we discuss the specific metrics used in our analysis.

#### Coverage

The first metric, which we call **Coverage**, represents what proportion of annotatable elements are actually annotated. Let *n* be the total number of annotatable elements, and *m* the number of elements with at least one annotation. The formula is defined as below:

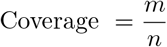

Since *n >* 0 and 0 ≤ *m* ≤ *n*, coverage is well defined and ranges between 0 and 1.

#### Consistency

Once a model element is annotated, next step should be to check whether its annotations are appropriate. **Consistency** indicates whether the annotation is appropriate, given the model element type. There are two questions to consider for each annotation term: i) Does the term exist within the corresponding knowledge resource? ii) Does the term match the type of the model element? For example, species should not be annotated with terms that represent chemical reactions.

We approached GO/SBO/CHEBI and KEGG/UNIPROT differently:

- GO/SBO/CHEBI: we first constructed a DAG for each knowledge resource, and checked if each term has a path to its appropriate root term. For example, GO:0000165 (MAPK cascade) will be a consistent annotation if it is annotated to a `libsbml.Reaction` instance, because there is a path from the term to the root GO:0008150 (biological process) and the root (GO:0008150) is considered appropriate for reactions. Alternatively, the terms associated with GO:0003674 (molecular function) are also considered appropriate for reactions. If the term did not exist within the ontology graph or was annotated to a wrong element type (such as `libsbml.Species`), the annotation would be considered inconsistent.
- UNIPROT/KEGG: since there is no hierarchy, we checked whether each ontology type matched the element type, and if the term existed within the current ontology database. The UNIPROT terms should be annotated only to `libsbml.Species`. For KEGG identifiers, we assumed KEGG Pathway and KEGG Reaction should be used for `libsbml.Model` and `libsbml.Reaction` instances, while KEGG Compound, KEGG Drug, KEGG Genes, and KEGG Orthology identifiers are suitable for `libsbml.Species` instances.

If a model element included more than one identifier, the element was considered consistent only if all identifiers to that element were consistent. To formulate this, let *M*_*i*_ be the set of annotations of *i*th annotated model element. Then the model-level consistency was calculated using the formula below:

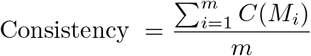

Where *C*(*M*_*i*_) ∈ {0, 1} such that *C*(*M*_*i*_) = 1 if all annotations for the *i*th element are consistent, and 0 otherwise. *m* is the number of annotated elements within a model.

The third and final metric is **Specificity**, which tests how precise (or detailed) the annotations are. Specificity is only calculated using model elements with consistent annotation. As in consistency, we used different methods for GO/SBO/CHEBI and KEGG/UNIPROT:

- GO/SBO/CHEBI: since the ontology terms were hierarchical, we used the concept of information content (IC) [9]. The information content of a term *t* is defined as:

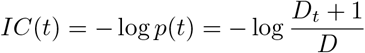

where *p*: 𝒞 → [0, 1], such that *p*(*t*) is the probability of encountering an instance of the term *t* in the set of possible ontology terms 𝒞 [15]. *D*_*t*_ is defined as the number of descendants of term *t*, and *D* is the number of all terms within the ontology. Using this information content, the specificity of a GO/SBO/CHEBI term is defined as the ratio of IC of the content to the IC of leaves in the tree, where leaves (*t*_*l*_) are defined as terms that have no children:

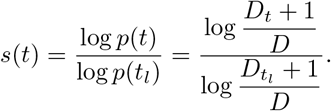
- UNIPROT/KEGG: since (i) there is no hierarchy between terms, and (ii) each term represents either an exact molecular compound or a chemical reaction, we assigned 1.0 to each consistent annotation term.

Combining all of the above, the specificity of a model is the average of *s*(*t*) from the consistent model elements:

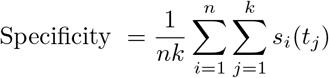

In the above formula, *t*_*j*_ indicates the *j*-th ontology term in the *i*-th consistent model element. If a model element contained more than one identifier from the same knowledge resource, they were averaged first.

We note in passing that, it may not be always the most desirable to annotate a certain element by only leaves, since the union of the information contained by all children terms may still be less than the information of their parent. However, in general, the biological concepts are considered more precise if they are deeper in the ontology tree [17].

The python package SBMate has implementation of the above three metrics and is provided as a `pip` installable package. It is hosted at https://github.com/woosubs/SBMate.

### 2.3 Extension of SBMate

SBMate was created so that the metrics can be easily added and extended by other modelers and developers. Currently, the `sbmate.AnnotationMetrics` class combines the coverage, consistency, and specificity scores into a `pandas.DataFrame`; one can create a new metric class and extend SBMate by modifying the keyword argument `metric_calculator_classes` of the `sbmate.AnnotationMetrics` class.

## 3 Results: Analysis of BioModels

The three SBMate can be obtained using the `sbmate.AnnotationMetrics.getMetrics()` method, with an SBML model file and specifying the output type, as shown below. The example below was executed on JupyterLab.

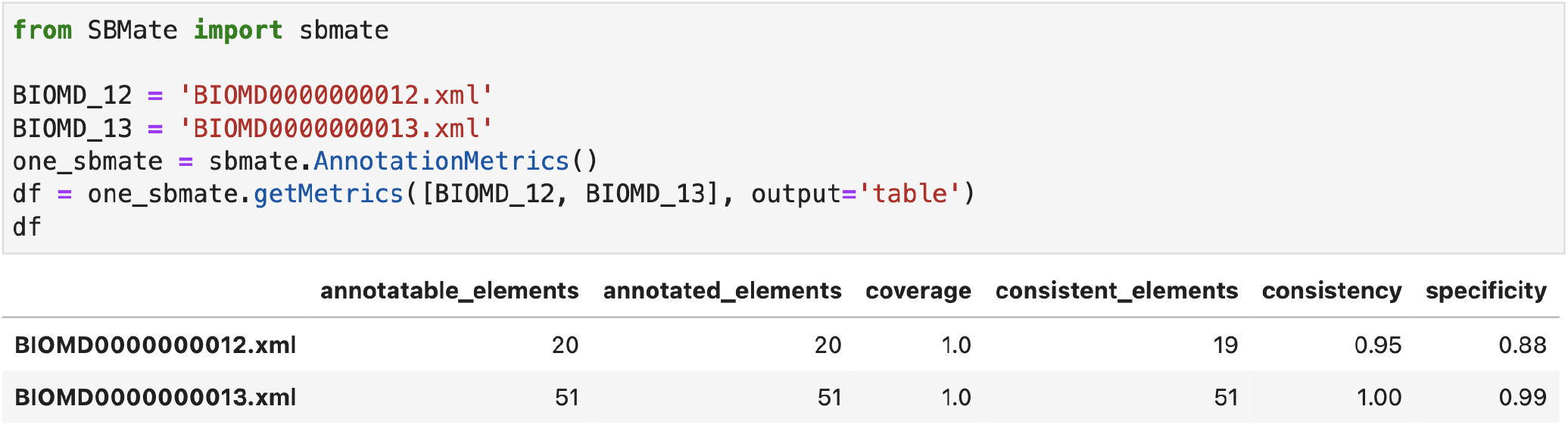

We analyzed 1,000 curated models from the BioModels repository (downloaded Mar 31, 2021), and results are displayed in Figure 3. Coverage score ranged between 0.0 and 1.0, where 145 models were fully annotated, while 88 models did not have any annotation for their annotatable elements. Except the models with such extreme values, coverage score seemed overall evenly distributed. Consistency score shows that, once a model was annotated, most annotataions tended to be consistent, as 477 out of 912 annotated models had consistency score 1.0, with only 4 models having zero consistency score. Specificity showed a similar pattern; 105 out of 908 models had specificity score 1.0 for their consistent elements and half of such models scored over 0.87. The lowest specificity score was 0.17. The statistics of the three metrics were summarized in Table 2.

**Table 2:** Summary of three annotation quality metrics. *Eligible Models* indicates the number of models where each metric could be applied. Only 912 models were checked for consistency as 88 models did not have any annotation in their annotatable elements. Specificity were calculated for 908 models, as 4 additional models did not have any consistent elements.

|  | Coverage | Consistency | Specificity |
| --- | --- | --- | --- |
| Eligible Models | 1,000 | 912 | 908 |
| Mean (Std) | 0.56 (0.34) | 0.93 (0.14) | 0.85 (0.14) |
| Median | 0.58 | 1.00 | 0.87 |
| Maximum | 1.00 | 1.00 | 1.00 |
| Minimum | 0.00 | 0.00 | 0.17 |

**Figure 3:**
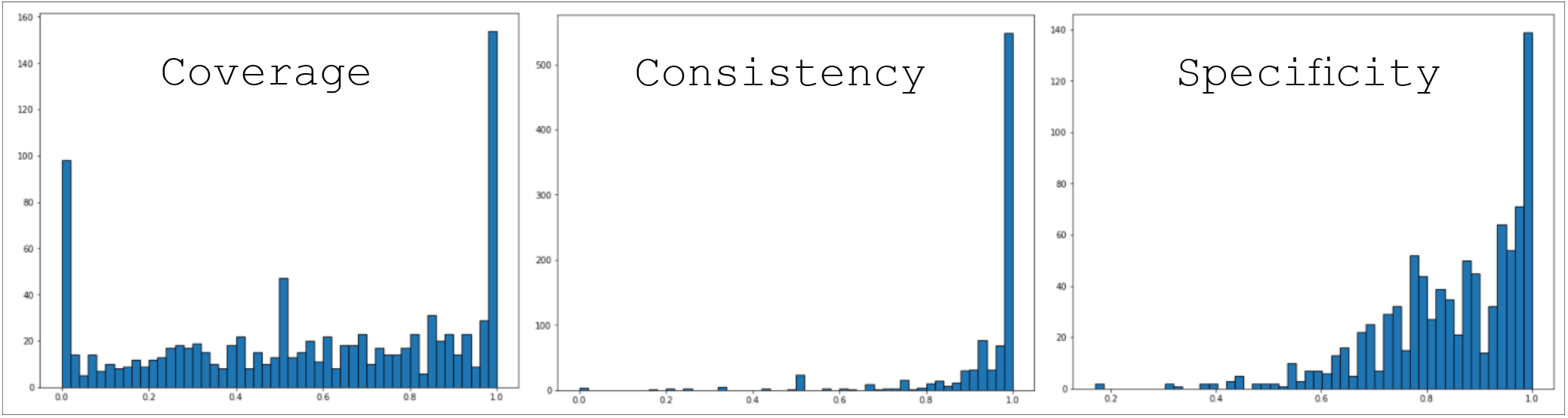
Distribution of annotations metrics of 1,000 BioModels. From the left, coverage, consistency, and specificity scores are displayed.

## 4 Conclusion

In the current version of SBMate, the three metrics are computed sequentially. First, the existence of annotation (coverage) is clearly a prerequisite for evaluating annotations. Once the model element is annotated with one or more identifiers, one can check whether the annotations are appropriate (consistency) for the type of model element and for the knowledge resource. Finally, the level of specificity of a model element can be measured if it is annotated with appropriate terms.

In our analysis of BioModels, we found that over 90% of all models in the repository contained at least some annotations. This might reflect the general agreement among researchers on the necessity of semantic annotations. However, most models were only partially annotated (i.e., some elements did not have annotation). Based on the consistency score, we found that once a model element was annotated, most of them were consistent with the type of corresponding model elements. Finally, the distribution of specificity score implies some of the current model annotations may need more tuning, as there were several models with low specificity scores, which may indicate lack of detail in such annotations.

Overall, we found the three metrics were able to describe the state of annotations in the existing systems biology models and to suggest proper ways to annotate new models. Our methods can also be used to upgrade existing annotations; for example, users can detect models with low specificity score, and replace the annotations with terms with higher precision. On the other hand, the current annotation quality scores may imply the difficulty of manual annotation by the modelers. This issue will become more prominent as the size and complexity of systems biology models grow. In future, we may need tools that assist model authors and curators annotate newly created models.

Our work could be improved in several ways. The current version of SBMate tests annotations in only four model elements: `libsbml.Model`, `libsbml.Compartment`, `libsbml.Species`, and `libsbml.Reaction`. Additional annotation checking on other elements, such as `libsbml.KineticLaw` and `libsbml.Parameter`, would help better describe the overall quality of annotations in the models. Second, we only used annotations that belong to the five knowledge resources: GO, SBO, CHEBI, UNIPROT, and KEGG. There are many other ontologies such as EC (Enzyme Commission)-Code and NCIT (National Cancer Institute Thesaurus), and SBMate could be extended to include such systems.

We have only considered models encoded in SBML but a wide range of models also exist in the CellML Format [3, 16]. Supporting CellML would require additional work because CellML stores models in pure mathematical form without any element information. In CellML, element information is provided purely in the form of annotations; the analysis would therefore be more involved. One way to mitigate this challenge is to analyze annotations stored as metadata files. For example, tools such as SemGen can annotate both CellML and SBML models and store the information in OMEX metadata files [12, 11]. We expect the future version of SBMate would include methods that can analyze OMEX annotation files so that both types of models can be analyzed on equal basis in spite of their structural difference.

The SBMate algorithms were established to provide a framework for evaluating annotations, and additional metrics can be easily added to extend the current version. We hope SBMate will be a useful tool to help model curators check the existing model annotations and model authors annotate new models.

## 5 Acknowledgements

This work was supported by National Institute of General Medical Sciences, National Institute for Biomedical Imaging and Bioengineering, and National Science Foundation awards P41GM109824, and 1933453, respectively. The content is solely the responsibility of the authors and does not necessarily represent the official views of the National Institutes of Health, National Science Foundation, the University of Washington, or the Center for Reproducible Biomedical Modeling.

